# Recruitment of Env to the HIV-1 T cell virological synapse by targeted and sustained Env recycling

**DOI:** 10.1101/2020.12.08.417188

**Authors:** Lili Wang, Alice Sandmeyer, Wolfgang Hübner, Hongru Li, Thomas Huser, Benjamin K. Chen

## Abstract

HIV-1 infection is enhanced by cell-cell adhesions between infected and uninfected T cells called virological synapses (VS). VS are initiated by the interactions of cell-surface HIV-1 envelope glycoprotein (Env) and CD4 on target cells and act as sites of viral assembly and viral transfer between cells. To study the process that recruits and retains HIV-1 Env at the VS, a replication-competent HIV-1 clone carrying an Env-sfGFP fusion protein was designed to enable live tracking of Env within infected cells. Using surface pulse-labeling of Env and fluorescence recovery after photobleaching (FRAP) studies, we observed targeted accumulation and sustained recycling of Env between the endocytic recycling compartment (ERC) and the VS. We observed dynamic exchange of Env at the VS while the viral structural protein, Gag, was largely immobile at the VS. The disparate exchange rates of Gag and Env at the synapse indicate that retention of Env is not likely to be maintained by entrapment into an immobile Gag lattice or through immobilizing interactions with CD4 on the target cell. A FRAP study of an Env endocytosis mutant showed that recycling is required for the rapid exchange of Env at the VS. We conclude that the mechanism of Env accumulation at the VS and incorporation into nascent particles involves continuous internalization and targeted secretion rather than irreversible interactions with the budding virus.

## INTRODUCTION

The HIV-1 envelope glycoprotein (Env) plays crucial roles as the surface glycoprotein on the virus particle, mediating virus binding, fusion and entry, as well as in initiating the formation of cell-cell adhesions that facilitate viral transmission, called virological synapses (VS) (1–3). HIV-1 can infect cells through cell-free virus, or through cell-to-cell routes which involve direct transfer of virus across a VS. HIV-1 Env is the surface antigen exposed on the surface of the cell or on virus particles where it can engages its main target CD4. HIV-1 is an enveloped virus that assembles and buds from the plasma membrane in a process mediated by the core structural protein Gag (4). An endocytic trafficking pathway helps to package Env into newly formed virus particles (5–7). The expression of Env at the cell surface renders infected cells susceptible to antibody detection, and while many antibodies against Env can block the formation of virological synapses, they are less efficient at blocking cell-to-cell infection than they are at blocking cell-free infection (8–12).

The biogenesis of HIV-1 Env begins at ribosomes on the rough endoplasmic reticulum (ER) where newly synthesized Env is glycosylated into precursor gp160 to form homotrimers (13). The cleavage of gp160 occurs in the Golgi apparatus by furin or furin-like proteases and results in two non-covalently associated peptides: a cell surface glycoprotein, gp120, and a transmembrane glycoprotein, gp41 (14, 15). Env trimers travel through the secretory pathway to reach the plasma membrane, and then are quickly recycled from the cell surface (16–20). This contributes to the very low number of Env glycoproteins on the cell surface. Lentivirus gp41 has a long intracytoplasmic C-terminal tail compared to other retroviruses (21). A membrane-proximal tyrosine-based sorting signal YxxL in the gp41 C-terminus interacts with the AP-2 to promote the internalization of Env (22–24). Env recycling from the cell surface to the endocytic recycling compartment (ERC) is a prerequisite for Env incorporation (6, 7). Proper incorporation of Env into viral particles also requires gp41 C-terminal sequences. The outward trafficking of Env from ERC to virus assembly area is mediated by C-terminal tyrosine-based motif YW795 (5).

HIV-1 cell-to-cell transmission leads to the efficient transfer of virus and infection (3, 10, 25) and mediates resistance to neutralization (8–12). Cell-to-cell transmission promotes viral diversity by supporting the co-transmission of multiple copies of HIV-1 per transmission event (26–28) and is proposed to play a role in escape from immune responses or may promote the evolution of drug resistance in settings of suboptimal therapy (9, 29, 30). The HIV-1 VS is an example of polarized viral transmission, where the assembly and release of Env and Gag are directed toward the receiving target cell, which internalizes the virus through an endocytic pathway (31, 32). At the VS, HIV-1 Gag, Env and CD4 localize to the site of cell-cell contact in an actin-dependent manner (3). Recruitment of Gag and Env protein and their transfer through VS occurs in a dynamic process following cell-adhesion (33, 34). Env-CD4 interaction is required for VS formation. Blocking the interaction of Env and CD4 with antibodies inhibits VS formation (10). During the formation of a VS, Env is observed to accumulate at the VS, however, the mechanisms of enrichment of Env at the VS are not well characterized. The extent to which Env diffuses laterally, is recruited to the VS from surface pools or may be concentrated by a secretory pathway that targets the VS is unclear.

The fusion of proteins with the green fluorescent protein enables live tracking of the protein within the cell (35). However, the relatively large size of GFP and its derivatives (30kD) requires careful consideration of the site of insertion to maintain the function of the protein of interest. Prior Env-GFP fusions have been expressed outside of the full proviral context or required complementation of WT Env to support viral replication (33). In order to preserve Env function, a strategy of insertion of GFP into the fourth variable loop was intended to yield a full-length infectious HIV clone with a functional Env (36).

Short peptide motifs in the Env cytoplasmic tail (CT) can control surface Env levels, direct incorporation of Env into viral particles, and can impact the conformation of the surface (36) domain of Env, which can further modulate Env fusogenic potential (37, 38). In this study, we engineered an infectious HIV-1 carrying a fluorescent-Env to observe the *de novo* expression of Env in an infected cell and track Env accumulation and turnover during VS formation. We followed the turnover rate of Env trafficking at the VS using fluorescence recovery after photobleaching (FRAP), which revealed that surface Env is constitutively recycled and the residence time at the cell surface is short lived measured in minutes, even at sites of high surface accumulation.

## RESULTS

### Engineering an infectious HIV carrying a sfGFP insertion into the Env V4 or V5 domains

To study the trafficking of Env to the VS we set out to design a fluorescent protein-tagged Env that is compatible with efficient packaging and viral membrane fusion. To minimize disruption of Env structural stability, we inserted a superfolder allele of GFP (39) directly into the HIV-1 Env coding sequences at selected points of V4 or V5 domain, which have previously been described as producing fluorescent Env (Fig.1A). The four HIV-1 clones carrying the Env-GFP fusion proteins produced similar levels of virus compared to the parent clone, HIV NL4-3 (Fig.1B). Three constructs produced virus with 25 to 50% of infectivity relative to HIV NL4-3 with a wild-type Env (Fig.1C). Western Blotting of cells producing HIV-1 Env-V4.1-sfGFP, HIV-1 Env-V4.2-sfGFP and HIV-1 Env-V5.2-sfGFP revealed the expected increase in size of the Env glycoprotein in the cell lysates as compared to WT Env from HIV-1 NL4-3 (Fig.1D). We noted that in cell lysates recombinant Env was processed to gp120-GFP fusion, but with a moderately lower efficiency. The recombinant Envs, Env-V4.1-sfGFP, Env-V4.2-sfGFP and Env-V5.2-sfGFP were also packaged efficiently onto virus particles. One recombinant construct Env-V5.3-sfGFP failed to produce full-sized Envelope proteins.

**Figure 1.**
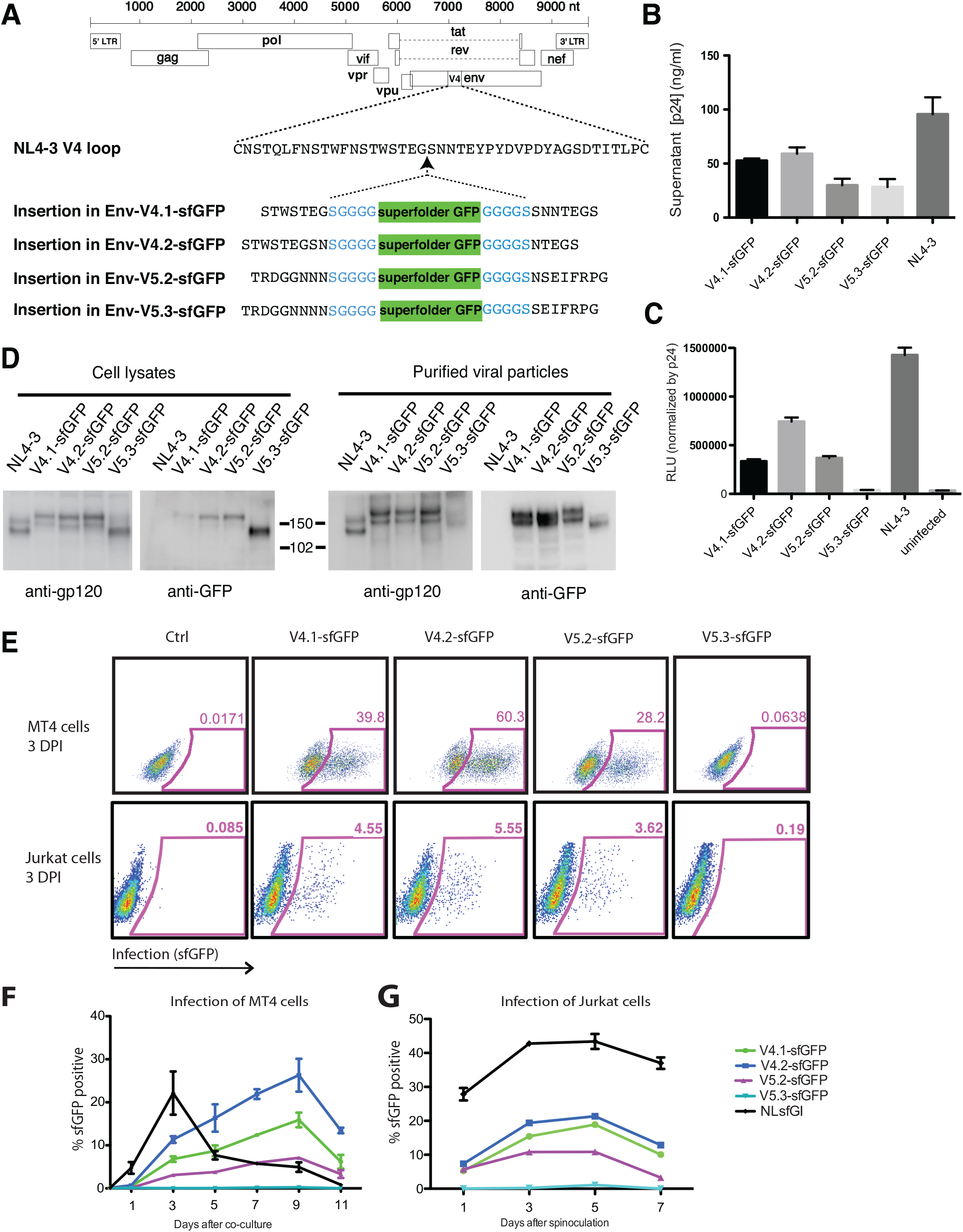
Construction of infectious HIV clones with fluorescent Env carrying sfGFP inserted into V4 or V5 domains of Env. (**A**) sfGFP is inserted into HIV-1(NL4-3) in V4 or V5. (**B**) Virus production by fluorescent Env HIV constructs following transfection of 293T cells. (**C**) Cell-free virus infectivity was tested by infection of indicator cell line, Tzm-bl. Tzm-bl cells were infected with supernatants with same amount of p24. (**D**) Western blot analysis of lysates of transfected 293T cells or of virus particles harvested from transfected cell supernatants and purified through a 20% sucrose cushion. Blots were probed with anti-gp120 or anti-GFP antibody. Viral supernatants and cell lysates were collected at 48 h post transfection. (**E**) Infection of Jurkat cells or MT4 cells with virus was assessed on day 3 after infection. (**F**) Infection of MT4 cells initiated by co-culture with HIV-nucleofected Jurkat T cells. FACS analysis was used to monitor the fraction of MT4 cells infected over time. (**G**) Infection of Jurkat cells initiated by co-culture with HIV-nucleofected Jurkat T cells. FACS analysis was used to monitor the fraction of Jurkat cells infected over time.

We next examined the efficiency of the four different HIV Env-sfGFP constructs to infect T cell lines. Infection of the highly permissive MT4 cell line was robust with cell-free virus showing high infectivity (Fig. 1E). Infection of Jurkat cells was lower in magnitude, with greater infection with HIV carrying V4.2-sfGFP followed by V4.1-sfGFP and V5.2-sfGFP (Fig. 1E). In both MT4 cells and in Jurkat cells, efficiency of infection of HIV V5.3-sfGFP was very low (Fig. 1E). To test if the four HIV clones carrying the Env-GFP fusion proteins can mediate spreading infection, Jurkat cells transfected with each of clones were co-cultured with MT4 cells or Jurkat cells. The spread of virus from transfected donor cells into target cells was measured using flow cytometry (Fig. 1F). The infection spread efficiently in MT4 cells with HIV-1 Env-V4.2-sfGFP replicating to a high peak titers as compared to wild type Env construct NL-sfGI, but with slower kinetics. In Jurkat cells, the HIV Env-V4/V5 sfGFP constructs all supported a spreading infection albeit with a lower efficiency compared with wild type Env construct NL-sfGI (Fig. 1G).

### Imaging HIV-1 carrying fluorescent Env constructs

To study the localization of HIV Gag and Env simultaneously during cell-to-cell spread of HIV-1, we created a series of three dual fluorescent HIV clones carrying a sfGFP fluorescent Env and a mCherry fluorescent Gag. We performed immunofluorescence staining of cells infected with HIV-1 Env V4.2 sfGFP-Gag-iCherry, carrying the Env V4.2 sfGFP, the chimeric Env which maintained highest infectivity, to compare the localization of V4.2-sfGFP Env to WT Env. Monoclonal antibody 2G12 binds to a non-conformational epitope and showed colocalization with Env-V4.2-sfGFP fluorescence in a sample cell (Fig. 2A-E). V4.2-sfGFP Env is abundantly expressed in cytoplasmic compartments, with the highest fluorescence shown in a peri-nuclear area, consistent with wild type Env distribution reported previously (16). To assess the distribution of Env and Gag relative to the plasma membrane, we performed structured illumination, super resolution imaging (Deltavision OMXv4.0 BLAZE) of Jurkat cells transfected with HIV-1 Env V4.2 sfGFP-Gag-iCherry and stained with a plasma membrane dye, Cell Mask deep Red (Fig. 2F-I). The predominant signal for Env was found in an intracellular compartment consistent with the trans-Golgi network (TGN), with minimal expression at the cell surface. A line projection of the fluorescence intensity across the plasma membrane revealed that Gag was located at the inner leaflet of plasma membrane. Env was not obviously enriched at the plasma membrane (Fig. 2F-J). Surface staining of Env on live cells expressing HIV Env-V4.2-sfGFP with anti-GFP antibody, showed puncta of Env at relatively low density (Fig. 2K-M). A time-lapse study of the kinetics of *de novo* expression of HIV Env-V4.2-sfGFP was performed using a confocal fluorescence imaging system from 6h to 26h post transfection (Fig. 2N). Env expression in the transfected cells peaked at 16-20h post transfection and declined thereafter (Supplemental Movie S1). Individual cells showed a similar peak expression of Env-sfGFP in the cells in the imaging field (Supplemental Fig. 1). To examine the distribution of HIV Env-V4.2-sfGFP during the formation of virological synapses, we co-cultured Env-V4.2-sfGFP transfected Jurkat cells with primary CD4^+^ target cells. Accumulation of Env at the junctions between HIV Env-V4.2-sfGFP transfected Jurkat cells and uninfected primary CD4+ T cells was observed (Fig. 2O-P). In primary CD4^+^ cells transduced with Env-V4.2-sfGFP viruses, a similar synaptic accumulation of Env was seen at the junction between the HIV-expressing primary T cell and the target primary T cell (Fig. 2Q).

**Figure 2.**
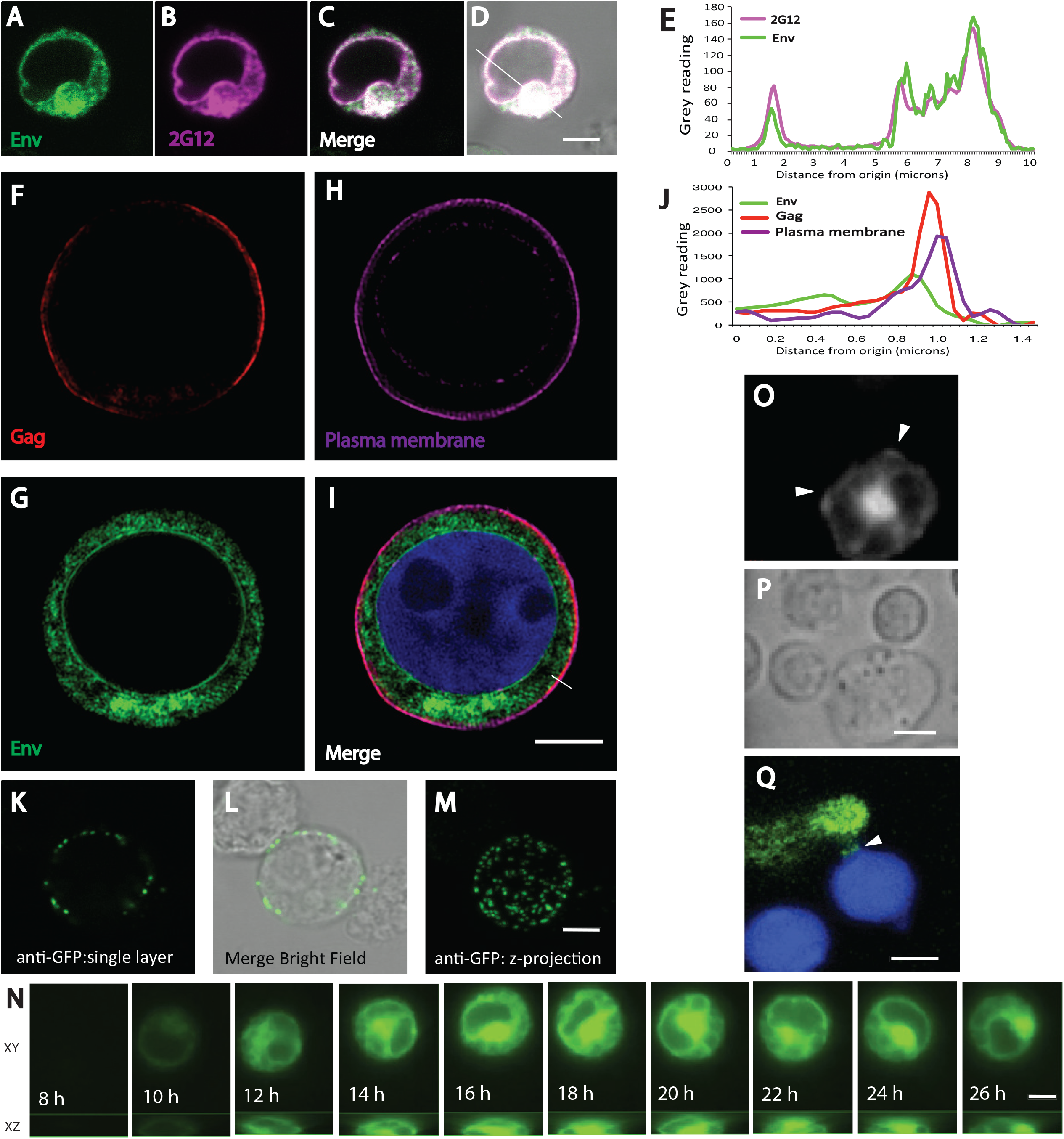
Fluorescence microscopy showing cellular distribution of sfGFP-tagged Env in Jurkat cells. (**A-D**) Confocal fluorescence microscopy imaging of Jurkat cells transfected with HIV Env V4.2-Gag-iCherry were fixed and stained with anti-Env mAb 2G12 (Magenta). **A**, V4.2 sfGFP Env localization, **B**, 2G12 Env immunostaining, **C**, Merged image, **D**, merged image with bright field overlay. (**E**) Graph shows the fluorescence intensity of Env and 2G12 staining traced along the line indicated in (**D**). (**F-I**) Super resolution structured illumination imaging of Jurkat cell transfected with V4.2-Gag-iCherry were stained with cell mask Deep Red. F, Cherry-Gag; G, sfGFP-Env, H, Cell Mask; I, Merged image. (**J**) Graph shows the fluorescence intensity of Gag, Env and plasma membrane along the line as indicated in (I). (**K-M**) Cell surface Env was stained with anti-GFP followed by secondary antibody while cells were alive at 4°C. The cells were then imaged for surface anti-GFP Env staining, K, single confocal plane; L, single plans merged with bright field; M, Z-projection of stack. (**N**) Confocal z stacks were acquired at 10-min intervals from 6 h post transfection for 20 h. Series of images show montage of fluorescence expression illustrating changes in the fluorescence pattern of Env-V4.2-sfGFP. (**O**) Env-sfGFP fluorescence is concentrated where two target cells make contact with a donor Jurkat cell. (**P**) is a bright field snap of (**O**). (**Q**) shows Env accumulation at VS area between an infected primary CD4 T cell and a target primary CD4 T cell. Bar: 5 μm.

### A dual fluorescent protein-expressing HIV with Gag-iCherry and Env-sfGFP participates in VS-mediated HIV transfer

HIV-1 constructs that carry a Cherry fluorescent protein inserted into Gag are not infectious, but generate highly fluorescent virus particles and participate in cell-to-cell transfer (10, 40). To determine if the fluorescent Env constructs are capable of participating in cell-to-cell HIV transfer across virological synapses, we generated dual fluorescent HIV which carry two fluorescent protein tags, Cherry and sfGFP, inserted into Gag and Env, respectively. The dual fluorescent viruses make abundant virus particles when transfected (Fig. 3A). The infectivity of these constructs in reporter cell lines are shown in Fig. 3B. These constructs maintained the ability to form VS and transfer Env and Gag into a target cell (Fig. 3C). HIV V4.2 sfGFP-Gag-iCherry expressing cells were tested for their ability to mediate HIV transfer across VS and transfer of fluorescent Gag and Env was observed (Fig. 3C). When the cell co-culture is treated with CD4 antibody, Leu3a, which can block CD4 engagement with Env, both Gag and Env transfer are blocked (Fig. 3C and D). Confocal fluorescence microscopy of the dual fluorescent constructs in Jurkat T cells and primary CD4+ T cells enabled visualization VS where both Env and Gag were colocalized (Fig. 3E, upper panel). In an example of a cell forming two virological synapses, one synapse showed both Gag and Env at the cell-cell junction, and the other showed accumulation of only Gag at the cell-cell junction (Fig. 3E, lower panel). During the imaging of virological synapses, Gag and Env colocalization at a virological synapse was more frequently observed soon after cell-cell mixing, and over time, the frequency of VS with only Gag concentrated at the VS increased. Images of VSs showed that Env and Gag were more frequently co-localized at 1 hour post coculture (82.4%); while after 3-hour coculture, the colocalization of Env and Gag at VS was observed in a lower percentage of cells (37.5%). Over time, both Gag and Env were observed to transfer into a target cell. The majority of fluorescent HIV proteins transferred into the target cells showed colocalization of Env and Gag, whereas some puncta appeared to represent the transfer of only Gag or only Env (Fig. 3G). Cotransfer of Env and Gag may be indicative of infectious virus, while the transfer of only Gag or only Env may represent the uptake of non-infectious viral antigen.

**Figure 3.**
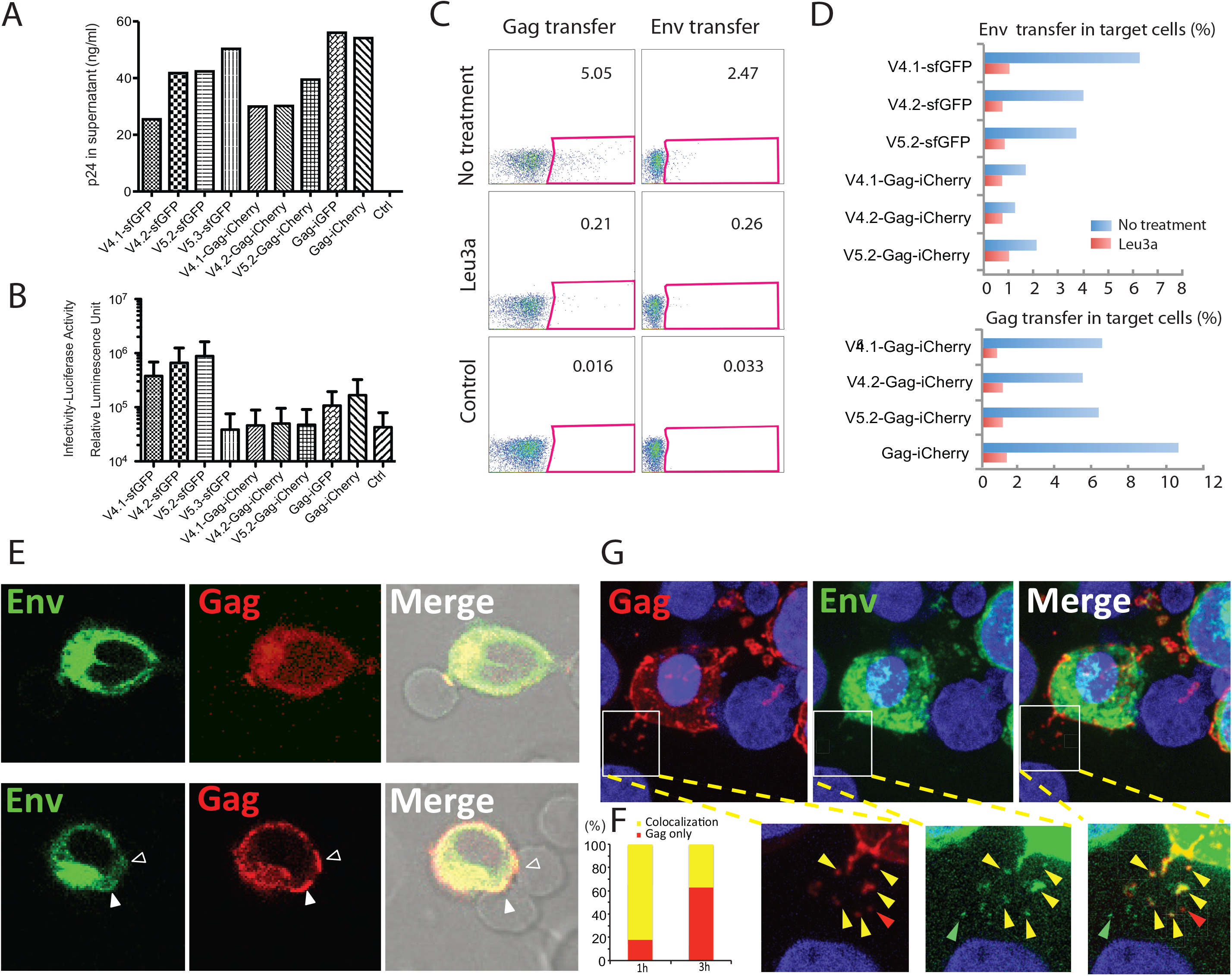
Cell-to-cell HIV-1 transfer assays using dual fluorescent construct of V4/V5-Gag-iCherry. (**A**) Dual fluorescent HIV-1 constructs produce viral particles in 293T cells as measured by p24 ELISA. (**B**) Infectivity of these dual fluorescent HIV-1 constructs using Tzm-bl assay shows infectivity of single fluorescent Env constructs and low infectivity of viruses carrying chimeric Gag-iCherry or Gag-iGFP. (**C**) Dual fluorescent constructs HIV-1 V4.2-Gag-iCherry participates in cell-to-cell transfer of HIV from Jurkat to primary CD4 T cells. Flow cytometry measures transfer of Gag-iCherry and Env V4.2-sfGFP signal following cell-cell co-culture, and the transfer is sensitive to CD4 antibody leu3a. (**D**) Cell-to-cell HIV-1 transfer of Gag and Env measured with indicated fluorescent HIV-1 constructs. (**E**) HIV-1 virological synapses between HIV V4.2-Gag-iCherry transfected Jurkat cells and primary CD4 T cells. Primary CD4 cells from healthy human blood were co-cultured with transfected donor cells for 3 h. upper panel: A typical synaptic button with both Gag and Env was shown between a donor Jurkat cell and a target cell. Lower panel: one donor cell nucleofected with V4.2-Gag-iCherry formed two virological synapses: the lower synapse shows both Env and Gag concentrated at the cell-cell contact site, while the upper synapse shows Gag accumulation without Env accumulation. (**F**) Analysis of Env and Gag colocalization at virological synapses. Samples fixed at 1 hour post co-culture and 3 hours post co-culture were compared. Virological synapses defined by Gag at the site of cell-cell contact were counted if Env was visible at cell contact site. (**G**) Transfer of both Gag and Env into target cells. Co-cultured cells were fixed and observed by confocal microscopy. Inset shows partial colocalization of transferred Gag and Env. Green, red and yellow arrowheads show Env only, Gag only transfer or co-transfer of both Gag and Env. Bar: 5 μm.

### Pulse-Chase labeling of surface Env tracks endocytosis and relocalization to the VS

The Env-CD4 interaction is a prerequisite of VS formation, but how Env is recruited to the VS is not clear. Prior imaging studies indicate that Gag is recruited from membrane associated pools and diffuses laterally into the VS (34). To examine the pathway of Env recruitment, we labeled cell surface Env with an anti-GFP fluorophore conjugated antibody and performed a pulse-chase imaging study to follow movements of surface-localized Env over time. Cell surface Env of a Jurkat cell nucleofected with HIV-1 V4.2-Gag-iCherry was visualized by staining at 4°C (Fig. 4A). Env is known to be quickly endocytosed from the cell surface (13). After warming cells to 37°C, cells were fixed after 5, 10 and 20 minutes to monitor the movement of pulse labeled Env. The surface Env stained cells were separated into two groups: one group that was mixed with target cells immediately after surface staining, and co-cultured at 37°C for 30 minutes. The second group was allowed to recover at 37°C for 30 minutes, then mixed with target cells for another 30 minutes. Both groups of cells were fixed afterwards and imaged with confocal microscopy. In group 1, surface labeled Env was mainly found in endocytic recycling compartments (ERC), while at the synapse area, no labeled Env was observed (Fig. 4C). In the second group, recycled surface Env localized mainly to the cell-cell junction when virological synapses were observed (Fig. 4D). These results indicate that surface-labeled Env can be endocytosed into the ERC, and then traffics specifically to the VS.

**Figure 4.**
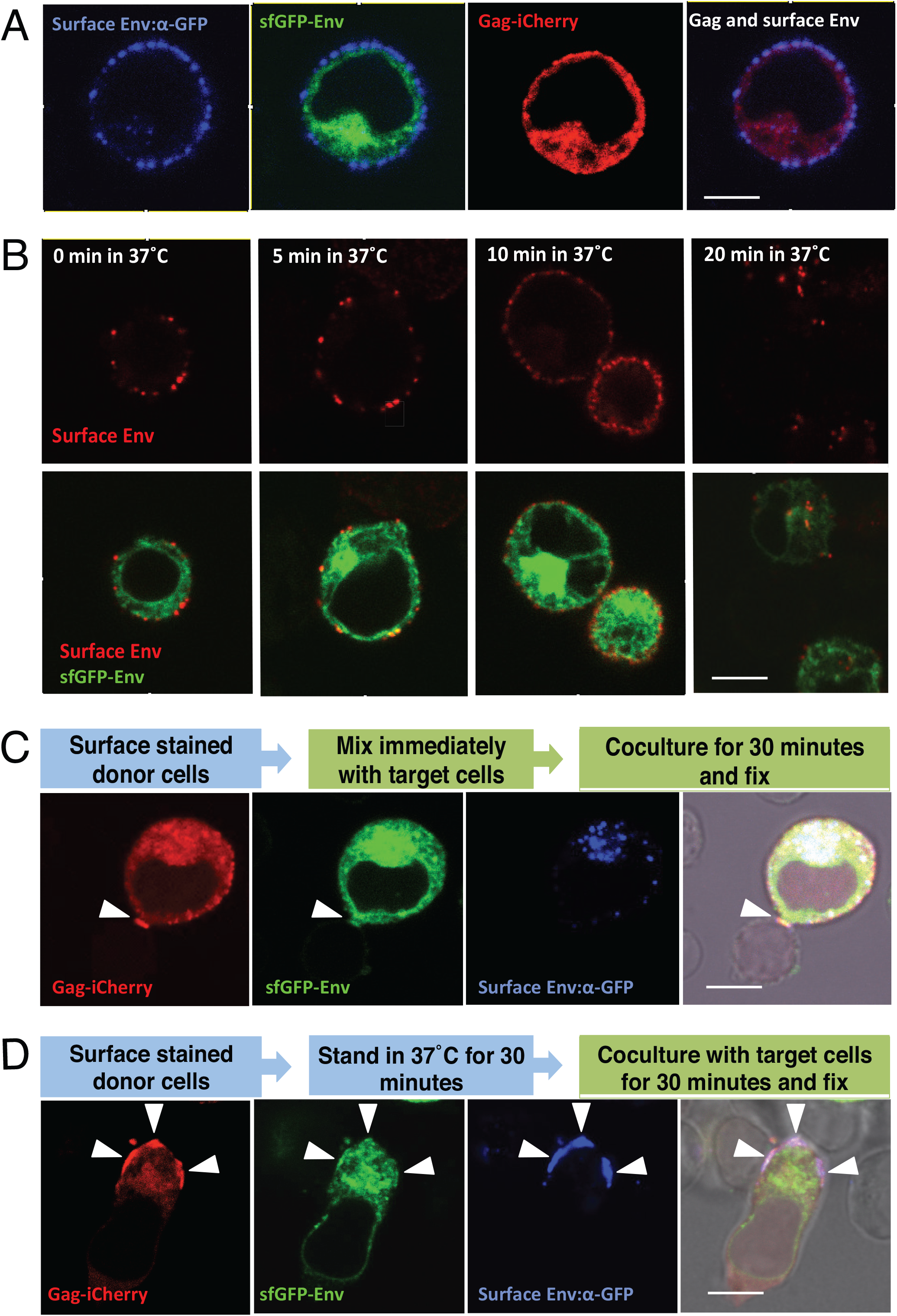
Pulse-chase labeling of cell-surface Env shows that recycled Env is targeted to virological synapse. (**A**) Live cell surface staining of V4.2-Gag-iCherry: nucleofected Jurkat cells stained with anti-GFP antibody at 4°C. (**B**) Pulse-chase of surface Env to determine time required for endocytosis: cells with surface stained Env were moved from 4°C to 37°C and kept for indicated time. The cells were fixed after incubation at 37°C and imaged. (**C**) The stained cells in (**A**) were immediately co-cultured with primary CD4 target cells for 30 min at 37 °C and fixed for imaging. (**D**) The stained cells in (**A**) were put in 37°C for 30 min first, then co-cultured with primary CD4 target cells for another 30 min at 37 °C and fixed for imaging. Arrowheads show virological synapses. Bar: 6 μm.

### Fluorescence recovery after photobleaching (FRAP) of HIV Env V4.2 sfGFP-Gag-iCherry at VS reveals constitutive turnover of Env at the VS

The results above indicate cell surface Env can be internalized into the ERC, reappear at cell surface, and then accumulate at the VS. How Gag recruitment may influence Env at the VS is not known. It is possible for instance that the recruitment of Gag to the VS may trap Env during its incorporation onto nascent virus particles, or that the interaction of Env with CD4 may immobilize it at the cell surface. To simultaneously track the kinetics of Env and Gag recruitment to the VS, we performed fluorescence recovery after photobleaching (FRAP) experiments with HIV V4.2 Env sfGFP-Gag-iCherry to measure the rate of turnover of Env and Gag at VS. We identified cells with a VS that showed both Gag and Env colocalized at the cell contact area. Half of the VS was photo-bleached, and the other half of the VS allowed segmentation of the VS and measurement of recovered fluorescence over time. Additional unbleached areas were tracked over time as a control to determine the basal rate of photodecay. With a smaller VS, the entire VS was bleached, and a nearby area was used as a control. As shown in Fig. 5A-1, the white square indicates the bleached area, and the yellow closed region is the selected region of interest (ROI). ROI-1 is the bleached synapse area, while ROI-2 is the unbleached control area. A steady recovery of Env intensity was observed within about 200 seconds, while in the same time period, there was minimal fluorescence recovery of Gag. Four additional FRAP studies on four different virological synapses were performed (Fig. 5A-2 to A-5, see Supplemental Movies 2-6). The recovery curve of Env was fitted to a one-phase exponential association function for each ROI (Fig. 5A-1 to A-5, right panels, ROI curves). The Env intensity before bleaching was set to 100%. The maximum recovery over the time frame of imaging was used to calculate an immobile fraction which differed between the different samples (Fig. 5B). In all the VS we observed, Gag fluorescence recovery was not observed, while Env fluorescence recovery occurred within 2-3 minutes with the half recovery time (Fig. 5C), indicating a much greater rate of Env turnover at the VS relative to Gag.

**Figure 5.**
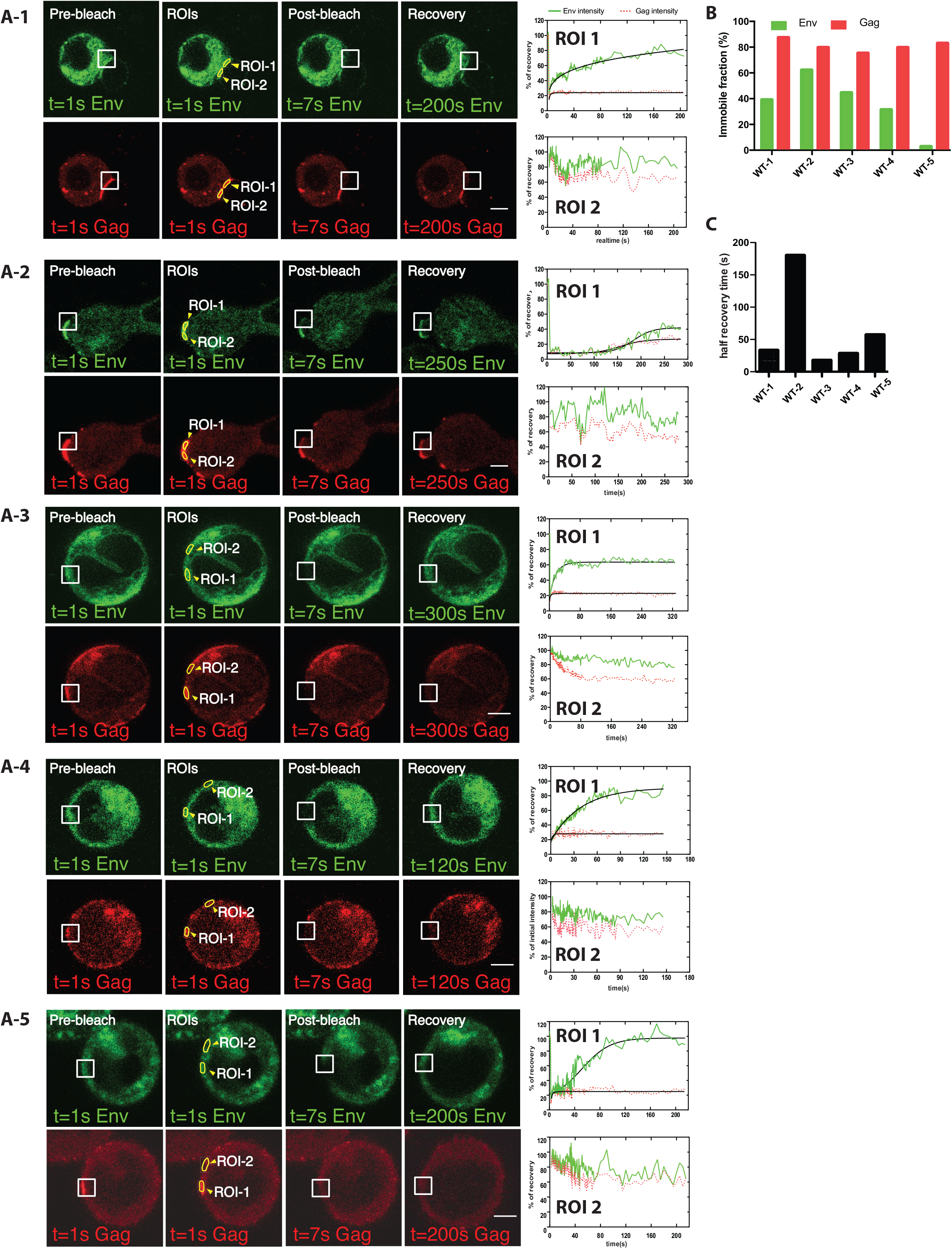
Rapid Env fluorescence recovery after photobleaching was observed at the VS. (**A**1) Before photobleaching a virological synapse with both Gag and Env could be observed between a donor cell and a target cell. A region covering part of the synaptic button is bleached as shown in the white square. After photobleaching, obvious fluorescent recovery was observed in Env, but not in Gag. ROIs were selected on bleached synapse or an unbleached area as shown in closed yellow region. A fluorescence intensity curve describing the fluorescence recovery is shown (left). Four additional representative cells repeating experiments with wild type V4.2-Gag-iCherry are displayed in (**A**2-**A**5). (**B**) shows the immobile fraction of each FRAP experiment. (C) shows the half recovery time of A1-A5. Bar: 3 μm.

### High turnover of Env at the VS requires endocytosis of Env using a membrane proximal tyrosine Y712

The gp41 C-terminal membrane-proximal tyrosine 712 in a YXXL AP-2 binding-motif is important for the internalization of surface Env through AP-2 mediated endocytosis (22). To test if surface Env endocytosis is required for synapse recruitment or turnover of Env at the VS, we introduced the Y712A point mutation into the viral clone, HIV Env-V4.2-sfGFP. HIV-1 with the Env Y712A mutation is reported to be less infectious as compared to wild type virus (38). We performed a T cell-to-T cell viral transfer assay using Jurkat donor cells and primary CD4 T cells as target cells and observed that cell-to-cell transfer of Env is increased by 3-fold in Y712A mutant relative to non-mutated virus in 3-hour co-culture (Fig. 6A). A separate cell-to-cell infection assay was performed to measure productive infection between HIV-expressing Jurkat cells and primary CD4 T cells. In this assay, both the wild type and the Y712A virus spread with similar efficiencies (Fig. 6B). In highly permissive MT4 cells, the Y712A virus spread with a slightly higher rate than wild type in 7-day productive infection (Fig. 6C). We next performed live imaging to see if the mutation which disrupts Env endocytosis from cell surface permits VS formation and accumulations of Env and Gag at the cell-cell junctions. We readily observed VS formation with high levels of Env recruitment to the synapse. When conducting FRAP studies we found that the Env recovery was dramatically decreased in the HIV-1 Env V4.2-Y712A-sfGFP when compared to the non-mutated clone (Fig. 6D-1). Four additional FRAP experiments were performed on virological synapses formed by HIV-V4.2-Y712A-sfGFP (Fig. 6D-2 to D-5). There was minimal or no recovery of Env or Gag observed over 5 minutes after photobleaching. Videos of all five virological synapses are in Supplemental Movies 7-11. Based on the extent of the fluorescence recovery the immobile fraction of Env was calculated, which was close to 100% in all the examples (Fig. 6E).

**Figure 6.**
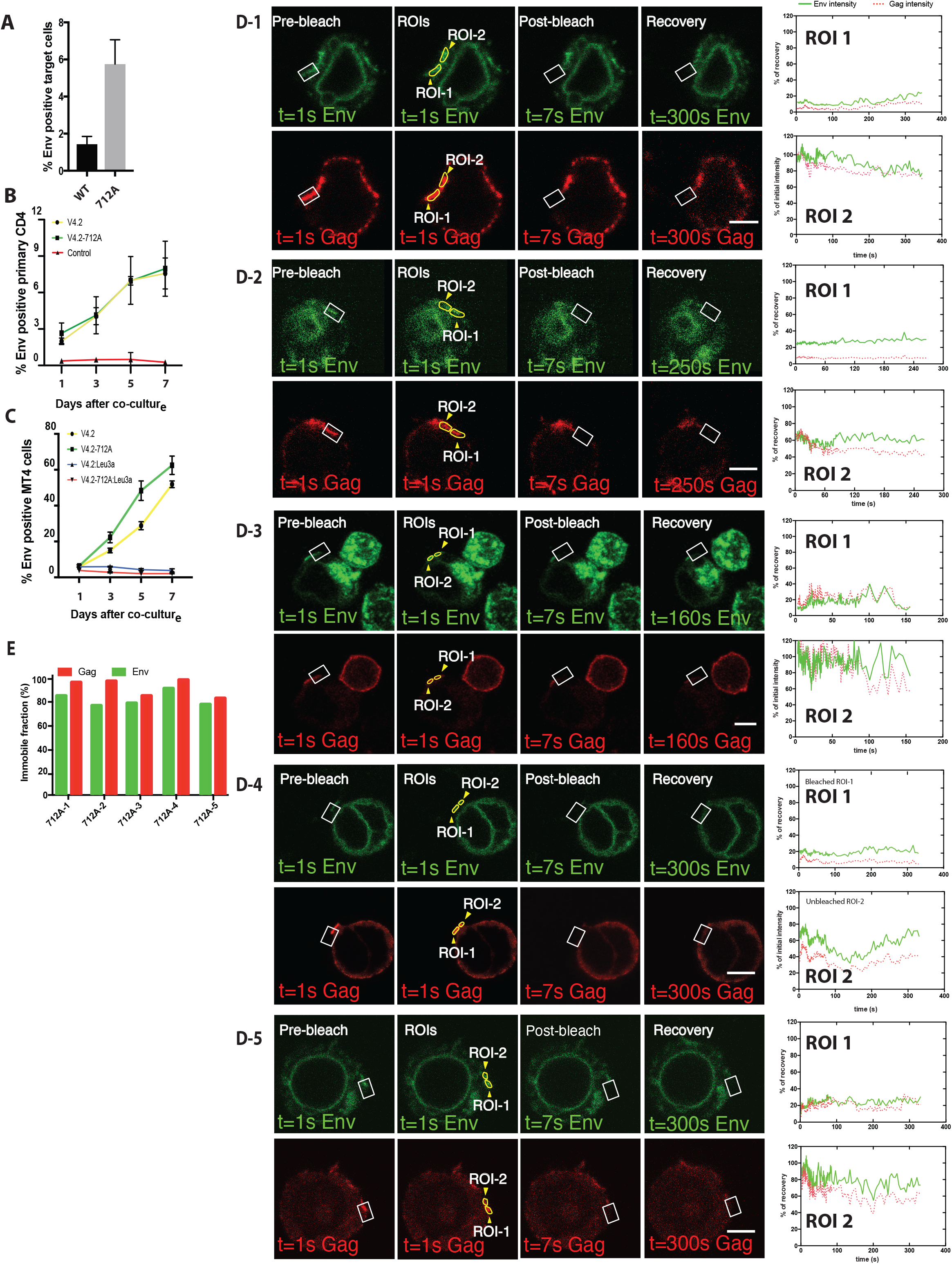
Env fluorescence after photobleaching does not recover when examining Y712A mutants of V4.2-sfGFP in FRAP. (**A**) Jurkat cells nucleofected with wild type Env-V4.2-sfGFP or Env-V4.2-Y712A-sfGFP were co-cultured with primary CD4 cells for 3 hours. Env transfer to primary CD4 cells were determined by Flow cytometry. (**B**) Jurkat cells nucleofected with wild type, Env-V4.2-sfGFP, or Env-V4.2-Y712A-sfGFP were co-cultured with activated primary CD4 cells to monitor productive infection in target cells. Samples were collected on day 1, 3, 5, 7 to determine the portion of primary CD4 cells with fluorescent Env. (**C**) Jurkat cells nucleofected with wild type, Env-V4.2-sfGFP, or Env-V4.2-Y712A-sfGFP were co-cultured with MT4 cells for days to monitor productive infection in target cells. Samples were collected on day 1, 3, 5, 7 to determine the portion of MT4 cells with fluorescent Env. (**D**) Fluorescence recovery after photobleaching (FRAP) of Env and Gag virological synapse with V4.2-712A-Gag-iCherry. Before photobleaching a virological synapse, both Gag and Env are concentrated at the junction between a donor cell and a target cell. A region of interest covering part of the synaptic button was bleached as shown in white square. ROIs were selected on bleached synapse (ROI-1) or an unbleached area (ROI-2) as shown in closed yellow region. Recovery curves of five individual experiments are displayed in (D1-D5). (**E**) shows the immobile fraction of each FRAP experiment. Bar: 5 μm.

## Discussion

In this study, we have constructed a fluorescent Env-carrying HIV clone that is capable of viral entry and productive infection in T cells in cell culture. The fluorescent Env fusion protein resembles wild type Env in its subcellular distribution and is very efficient in its ability to participate in VS formation and cell-to-cell infection. This is the first description of a HIV-1 clone encoding a fluorescent Env that is autonomously infectious. It enables live tracking of Env and its exchange between subcellular compartments during its recruitment to the VS. This tool makes it possible to observe Env distribution and trafficking within the context of productive infections, and in the absence of helper virus. We employ it here to test models for how Env trafficking contributes to viral spread between cells and supports the production of infectious virus particles.

Immunofluorescence with monoclonal antibody, 2G12, which recognizes a carbohydrate epitope, revealed that the localization of V4.2 Env resembles native HIV-1 Env. With super-resolution imaging and surface Env staining, we observed that the majority of Env is expressed in internal compartments, the endoplasmic reticulum, the Golgi apparatus, and endosomal compartments. As previously appreciated, cell surface Env represents a small fraction of total Env in the cell, and the results of our fluorescence microscopy also show very low surface Env levels (41–43), which also appears to correlate with the low Env density on viral particles (7-14 Env trimer/particle) (44, 45). When imaging VS, the fluorescent Env construct revealed increased concentrations of surface-targeted Env at cell-cell contact zones. This localization is consistent with the earliest VS imaging studies on fixed samples that indicate that Env accumulates to the VS area through actin dependent processes (3). In our study, cell surface Env that was not localized to the VS was only readily observed after amplification with fluorescent secondary antibodies. When visualized with GFP alone, V4.2 Env density at the cell surface was relatively sparse and evenly distributed, with no obvious areas where Env is pre-accumulated prior to VS formation (Fig.2 K-M and Fig.4 A).

The Env distribution before and after VS formation exhibits two different patterns diffuse versus focal. These patterns may represent different secretory pathways that can be polarized to traffic when cells are engaged in immunological synapses (46–48). The initial broad distribution of Env on the cell surface occurs prior to target cell engagement, and retargeting of the recycled Env to the VS appears to occur following CD4 engagement and may facilitate efficient particle incorporation. Evidence of VS-targeted Env trafficking can be observed prior to accumulation of Gag at the VS. When an infected cell is attached to an uninfected target cell, Env accumulation can be observed within minutes after cell attachment (33).

To explore the relationship of Gag and Env during the formation of VSs, a dual-fluorescent virus carrying Gag-iCherry and Env-V4/V5-isfGFP fusion proteins was studied. The dual fluorescent construct can also efficiently engage in cell-to-cell transfer of HIV-1. The ability to mediate cell-to-cell HIV transfer indicates that the CD4 binding sites of these constructs are fully functional, and signaling events prior to and during VS formation are intact. Using this construct, a surface labeled Env pulse-chase experiment indicated that the display of Env on cell surface is followed by internalization and subsequent concentration at the VS. In cases where an Env:CD4 dependent adhesion was formed between an infected and uninfected T cell, the labelled Env appeared to be directionally targeted to the cell-cell contact site with minimal signal observed away from the VS (Figure 4D). We suggest that Env functions initially as a cell-adhesion molecule and “detector” of target cell engagement, and then subsequently signaling from the cell-cell adhesion determines the site of polarized egress.

Early confocal imaging studies revealed the VS as a site where button-shaped accumulation of Gag formed at the adhesive junction between an infected cell and a target cell (34). Electron microscopy of the virological synapse revealed Gag accumulation in electron dense crescents forming a tight lattice at the VS (32, 34). Recruitment of Gag to the VS occurs from the lateral migration of plasma membrane-targeted Gag that moves towards the site of cell-cell contact site over minutes (34). FRAP studies here show that at a late stage of VS formation, after Gag synaptic button is established, Gag is largely immobile, and shows no recovery after photobleaching at the VS. This consistent with a largely irreversible incorporation of Gag into nascent budding particles (49). Compared to Gag, Env can also be observed at cell-cell contact area but at a lower relative concentration (Fig. 3F). A proposed model for Env incorporation into a budding virus particle is that it may be mediated by “trapping” of Env with its long cytoplasmic tail becoming encumbered in the 2-dimensional Gag lattice (50). However, in contrast to Gag at the VS, which is not exchanging with other pools of Gag in the cell, a large majority of Env continues to exchange with intracellular pools even after stable VS formation. This ability to exchange freely may indicate that a large fraction of Env is incorporated after Gag crescent formation at a late stage of assembly, where is does not get encumbered by the budding Gag lattice. This could be consistent with a recent superresolution imaging study suggests that Env is packaged at a late-stage of assembly and is localized with a distribution biased toward the necks of budding viruses (51).

In our FRAP studies (Fig. 5), the majority of Env at the bleached area recovered within minutes of photobleaching with some minor differences in the final immobile fraction for Env. This indicates the while most Env is continuously recycling to the VS a relatively small, variable fraction can be immobilized at the VS. The state of the cell, the stage of VS formation and the size of the VS all may contribute to these differences in the immobile fraction. The biosynthesis of HIV Env and Gag occur through different pathways. In this paper, our FRAP studies indicate that the forces that maintain Gag and Env at the VS are distinct. They reveal that physically they are not part of a stable complex during VS formation. The high degree of recovery after FRAP also indicate that a majority of Env is not held in place at the VS by the interaction with CD4 on the target cell. The results also indicate that the interaction between Env and CD4 at VS is reversible and mediated by a state that does not yet trigger viral membrane fusion.

We characterized an endocytic Env mutant and performed FRAP at the VS and observed that Env could still accumulate at the VS, however, the recycling of Env to the VS was not observed. This shows that blocking the endocytosis of Env with a Y712A mutation abolishes the turnover of Env at the VS. In this case, the accumulation of Y712A Env at the VS may be driven by the high concentration of Env at the cell surface. Truncation mutants in the C-terminal tail of Env or elimination of the main endocytic motif, Y712, allow high levels of Env to be displayed on cell surface (20, 24). An intact cytoplasmic tail is required for incorporation into the “neck” of the emerging budding virus and it is suggested that Env that are missing the CT are passively incorporated into viral particles (51). In our experiments, the Y712A endocytosis mutant leads to more viral transfer through the VS, though shows limited impact on the overall infectivity. This mutant can display different phenotypes depending upon the cell line it is tested in though in general it is still infectious (52). Together these data indicate that recycling is dispensable for VS formation, transfer and infection. We therefore speculate that a major role of recycling of Env at the VS lies in immune evasion: keeping surface Env density low to escape from immune surveillance (53) (54). Other studies from our group have shown Y712A mutants can also impact Env cell surface conformation and modulate the ability of broadly neutralizing monoclonal antibodies to neutralize cell-to-cell infection (55).

In summary these imaging studies support an emerging model of HIV-1 cell-to-cell infection, where Env traffics between the cell surface and the ERC before being packaged onto a budding virus particle (Fig. 7). An initial transient phase of exposure at the cell surface participates in the detection of the target cell. Subsequently Env that is recycled from surface to ERC, is redirected specifically to the VS, where Env is incorporated into virus. Dynamic trafficking of Env supports initial VS formation and enables VS to form under conditions where surface Env concentrations are maintained at very low levels. The process of recruitment to the VS is therefore optimized to reduce promote efficient transfer of virus from cell to cell while maintaining minimal surface expression of the dominant viral surface antigen.

**Figure 7.**
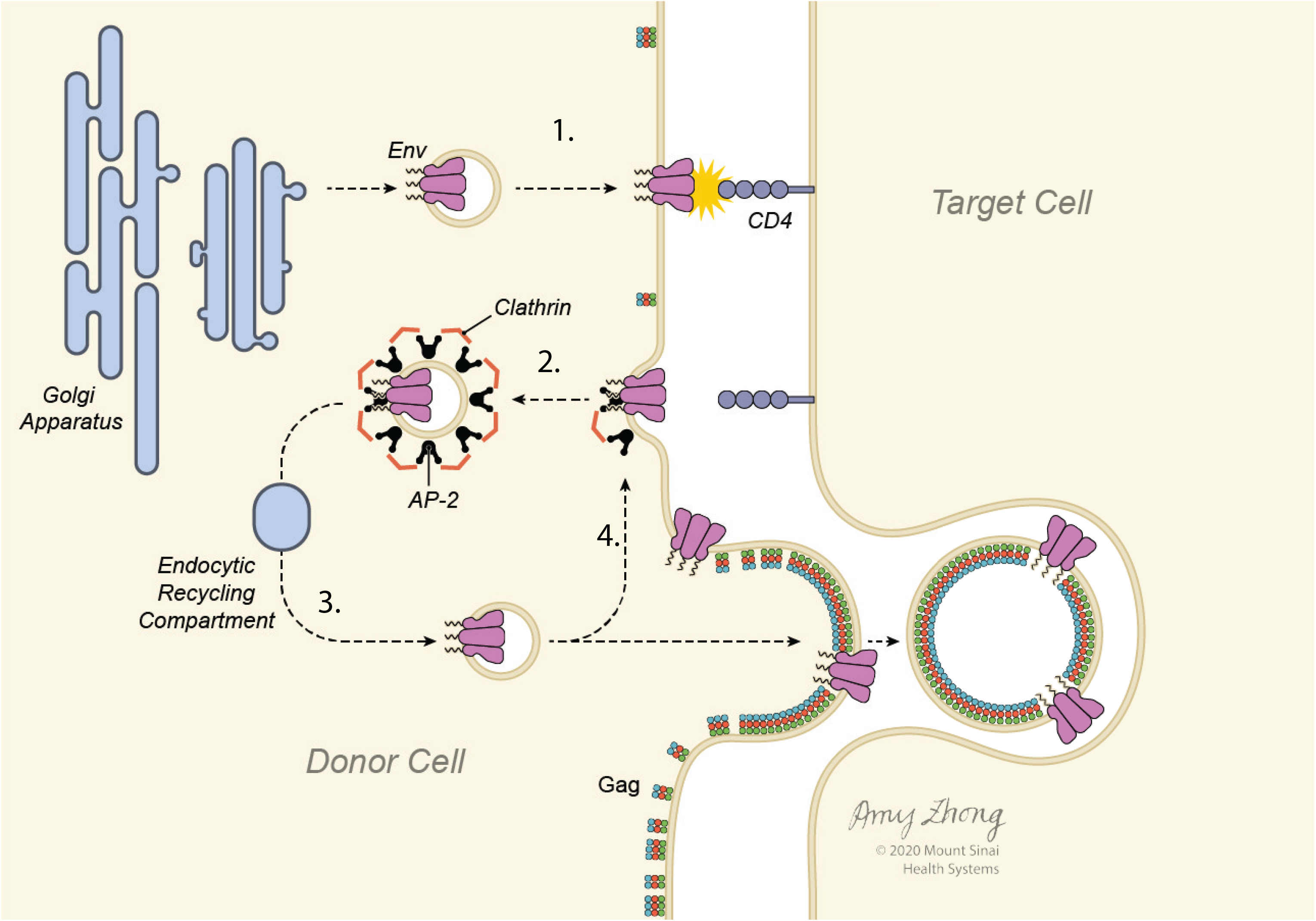
Model of Env trafficking pathways that support Env accumulation at the VS. (**1**) Env is transported to the cell surface following synthesis through the ER/Golgi pathways. (**2**) Clathrin-mediated endocytosis is initiated by recognition of the Env cytoplasmic tail by adapter protein complex, AP-2, which recognizes the membrane proximal tyrosine motif in Env. (**3**) Following internalization Env is recycled back to the cell surface, selectively trafficking to the VS where it can be incorporated into nascent virus particles. (**4**) Env at the VS continues to recycle while Gag does not exchange.

## METHODS

### KEY RESOURCES TABLE

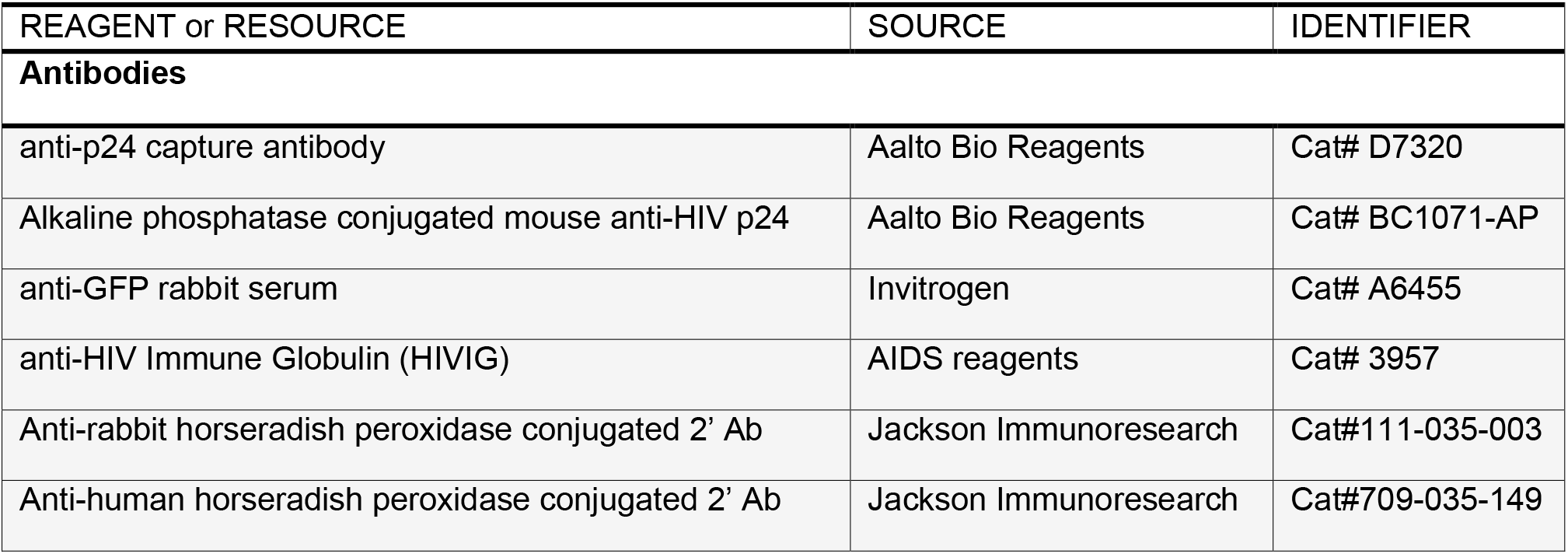
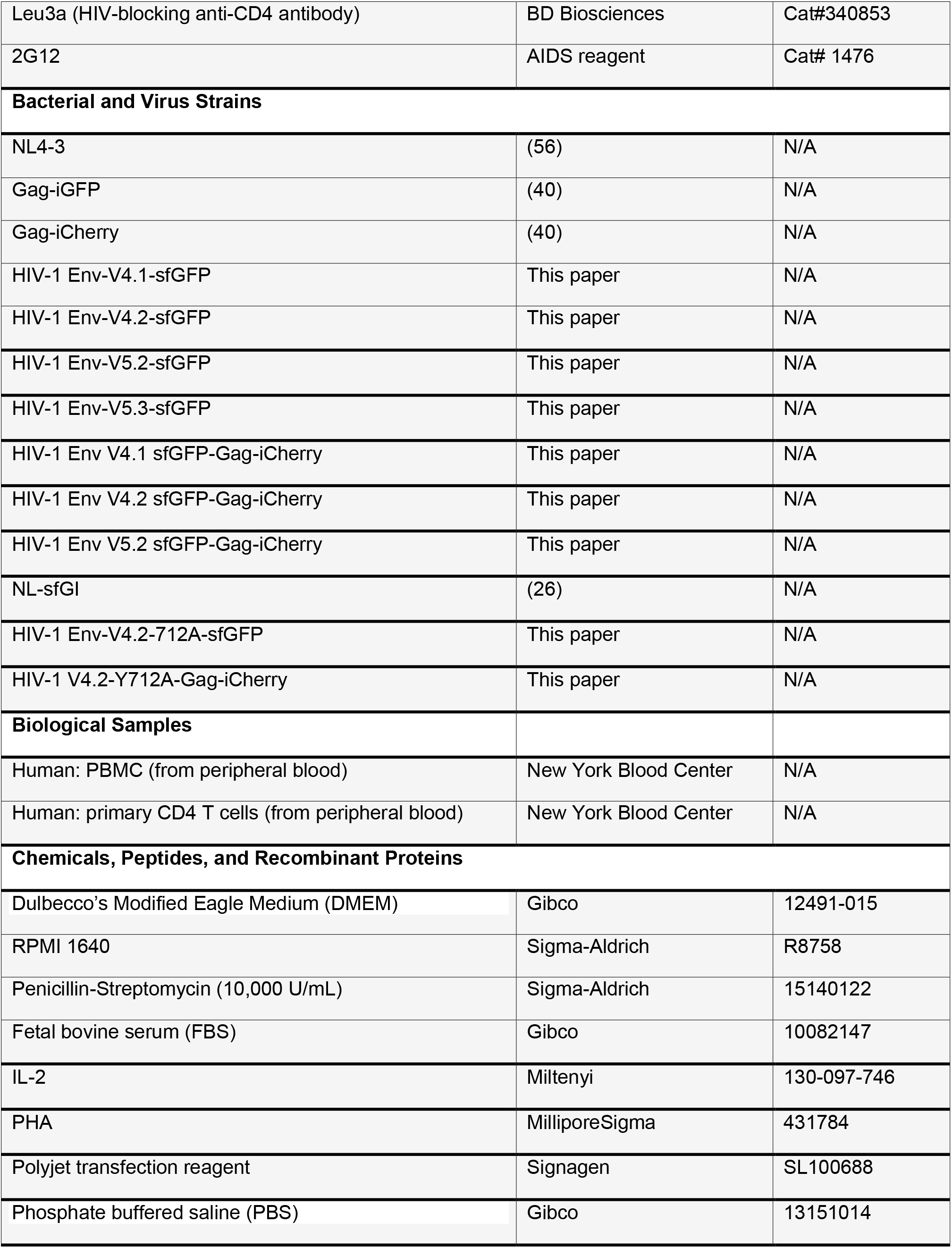
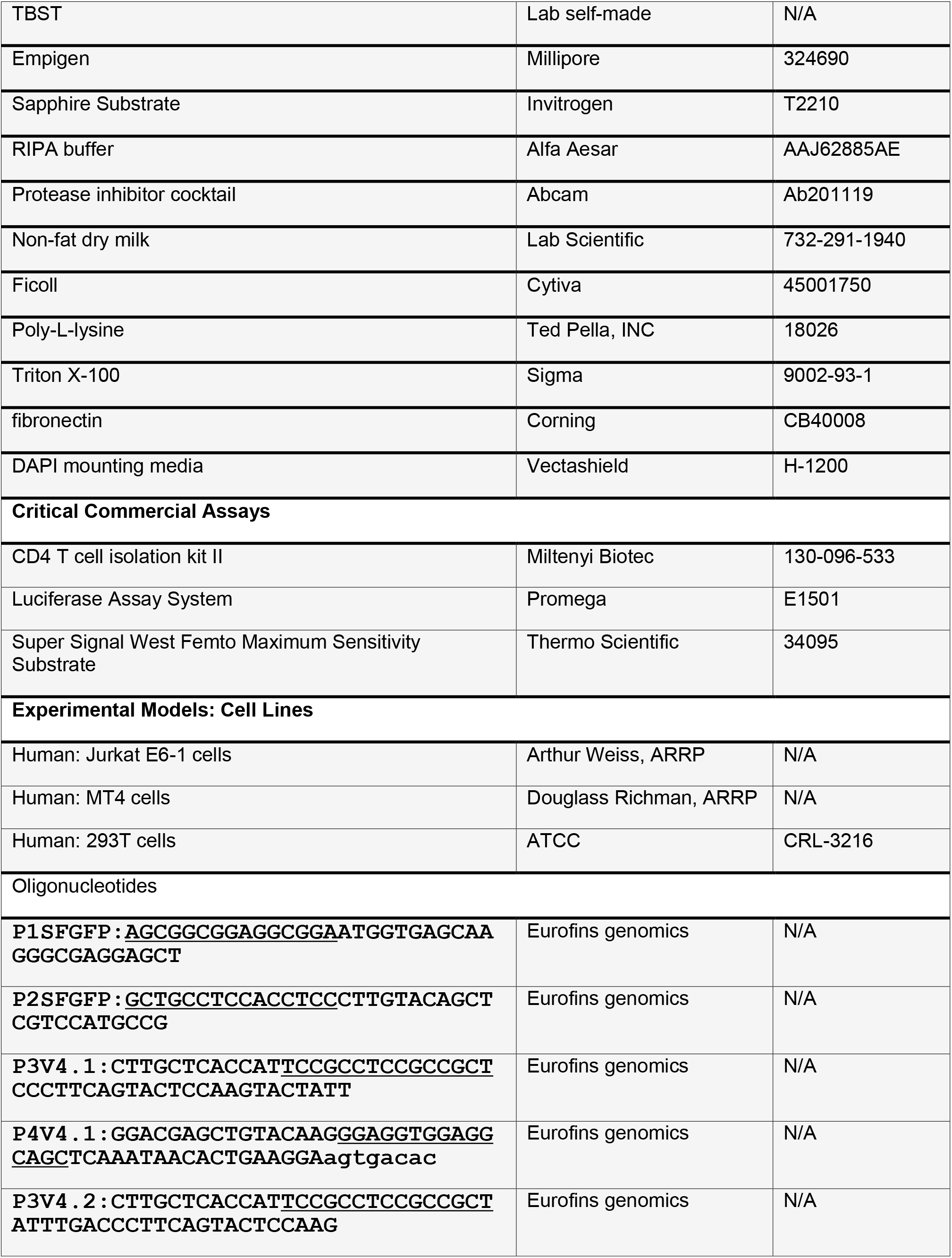
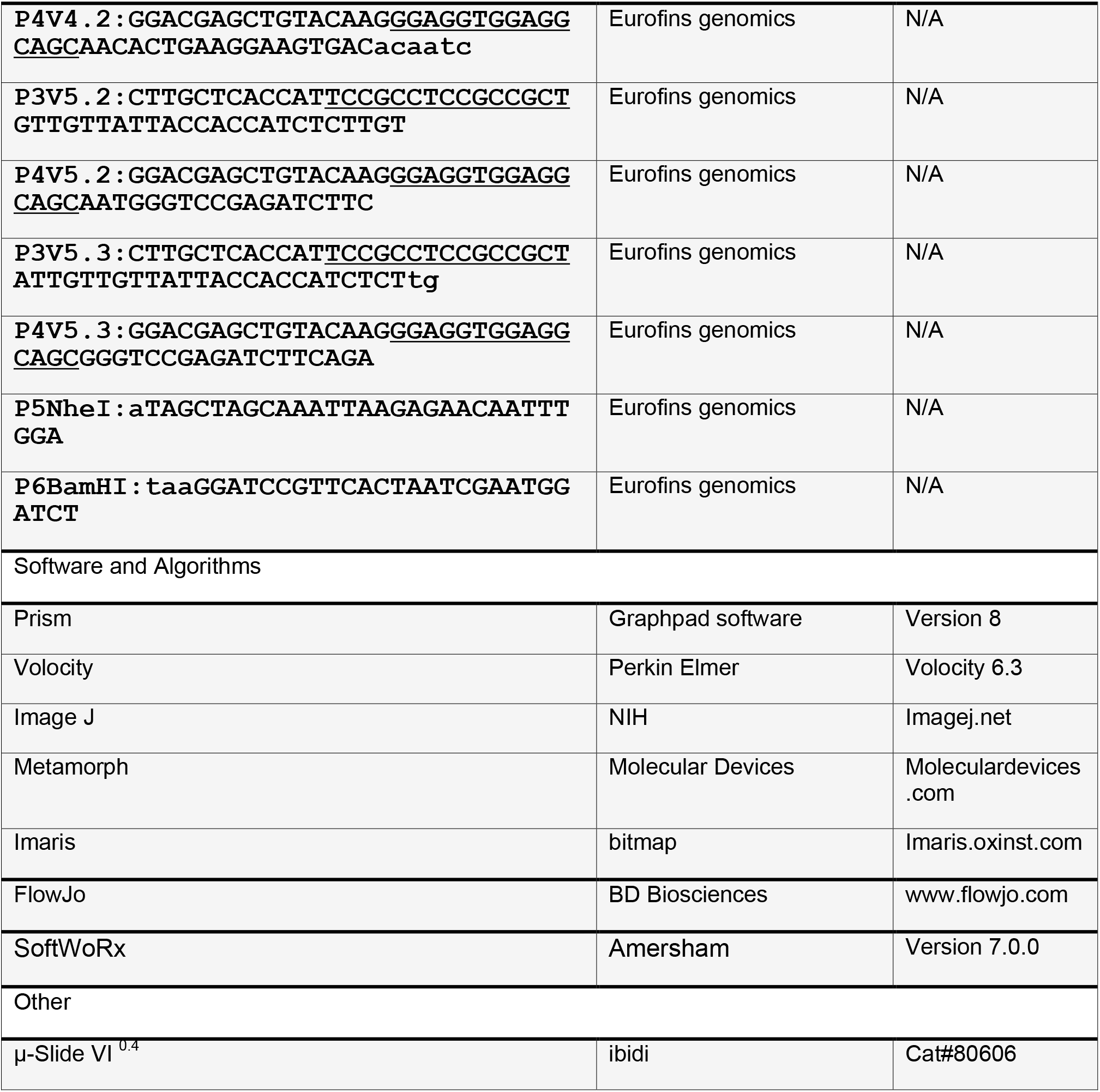

### CONTACT FOR REAGENT AND RESOURCE SHARING

Further information and requests for resources and reagents should be directed to and will be fulfilled by the Lead Contact, Benjamin K. Chen. Distribution of fluorescent Env HIV lab strains will require signing Material Transfer Agreement (MTA) in accordance with policies of Mount Sinai Medical Center.

### EXPERIMENTAL MODEL AND SUBJECT DETAILS

#### Cell lines

The CD4^+^ T-cell line Jurkat CE6.1 (ATCC) and CD4^+^ T-cell line MT4 were maintained in RPMI 1640 with 100 U/ml penicillin, 100 U/ml streptomycin and 10% fetal bovine serum (FBS). Cells were maintained at concentrations of less than 10^6^/ml. Primary CD4^+^ T cells were obtained from human peripheral blood from deidentified HIV-negative blood donors, through the New York Blood Center and CD4+ cells isolated by negative selection with a Miltenyi CD4 T cell isolation kit II (Miltenyi Biotec). Cell-free virus was produced by transfection of 293T cells in 10 cm plates using polyjet (Signagen). Media was exchanged 16h post transfection and virus supernatants were harvested 48h post transfection.

#### Human primary CD4 T cells

Human primary CD4^+^ T cells are obtained from peripheral blood with CD4 T cell isolation kit II (Miltenyi Biotec). Unactivated CD4^+^ T cells were maintained in complete RPMI medium containing 50 U/ml interleukin 2 (IL-2; ARP). Activated primary CD4^+^ cells were induced by co-culture with radiated PBMC feeder cells plus 100 U/ml IL-2 and 4 μg/ml PHA for 3 days.

#### Viruses

HIV Gag-iGFP and HIV Gag-iCherry are full-length molecular clones of HIV based on NL4-3 (Adachi et al.) previously designed to carry the green fluorescent protein (GFP) or mCherry protein inserted between the Gag MA and CA domains (40). HIV constructs with fluorescent Env were constructed by inserting Superfolder green fluorescent protein (sfGFP) internally into the Env V4 or V5 domains, designated HIV Env-HIV V4.1-sfGFP, HIV Env-V4.2-sfGFP, HIV Env-V5.2-sfGFP or HIV Env-V5.3-sfGFP. The superfolder GFP is introduced by 2-step PCR with the primers shown in key resource table. These fluorescent Env genes are also inserted into the context of HIV Gag-iCherry to yield constructs carrying Gag-iCherry and Env-sfGFP in cis. Y712A mutant was introduced by site mutation primer shown in key resource table.

### METHOD DETAILS

#### p24 ELISA

Costar 3922 flat-bottomed, high binding plates were coated with anti-p24 capture antibody overnight (Aalto D7320; 1:200 in 0.1M NaHCO3). Plate was washed twice with 1x TBST and blocked with 2% nonfat dry milk (Lab Scientific) for 1h then washed in TBST. HIV supernatants treated with 1% Empigen (1:100 and 1:1,000 in DMEM) along with titration of p24 standard are added to wells and incubated at room temperature for 2 hours, then washed 4x with TBST. Alkaline phosphatase conjugated mouse anti-HIV p24 (CLINIQA) was added (1:8,000 in TBST 20% sheep serum) and incubated for 1 hr followed by 6 TBST washes. 50μl of Sapphire Substrate (Tropix) was added to each well and incubated for 20 minutes. Luminescence was quantitated on Fluo Star Optima plate reader and sample values calculated based on nonlinear regression of standard curve using Prism software (Graphpad Inc.).

#### Western Blot Analysis

Cells or virus were lysed with RIPA buffer and protease inhibitor cocktail (Sigma). Protein loaded from viral lysates were normalized to p24 antigen content. Lysate equivalent of approximately 2×10^5^ cells per well were run on NuPage 4-12% Bis-Tris Gel (Novex) and transferred to Amersham Hybond-P PVDF membranes (GE Healthcare). Membranes were blocked with 2% nonfat dry milk (Lab Scientific), then probed with rabbit anti-GFP serum (1:5,000) or human anti-HIV serum (1:10,000) primary antibodies followed by anti-rabbit (Jackson Immunoresearch) or anti-human horseradish peroxidase (Jackson Immunoresearch) conjugated secondary antibody. Detection of band is using Super Signal West Femto Maximum Sensitivity Substrate (Thermo Scientific).

#### TZM-bl assay

Cell-free viruses were produced in 293T cells. TZM-bl cells were plated at 2×10^4^ cells/well in 96-well plates and incubated at 37°C with indicated viruses. Media was replaced after 24h of infection and incubated for another 24h. At 48h post infection, Media was aspirated followed by lysis in Luciferase Cell Culture Lysis Reagent (Promega). 20μl of each sample was read on Fluo Star Optima plate reader with injection of 50 μl of Luciferase Assay Reagent (Promega).

#### Cell-to-cell transfer assay

HIV-1 proviral constructs were transduced into Jurkat cells (donor cells) using Amaxa nucleofection as previously described (Amaxa Biosystems). In brief, 5 μg of endotoxin-free HIV-1 proviral plasmids was nucleofected into 6×106 Jurkat cells using Cell Line Nucleofector kit V, program S-18. Twenty hours after nucleofection, viable Jurkat cells were purified by centrifugation on a Ficoll-Hypaque density gradient, washed with complete buffer, and recovered at 37°C for co-culture. Unactivated primary CD4+ T cells (target cells) were cultured overnight in complete RPMI medium containing 50 U/ml IL-2. Donor and target cells were mixed at a ratio of approximately 1:1 and cocultured at 37°C for 3 h before they were treated with trypsin and fixed. Where inhibitor Leu3a, an HIV-blocking anti-CD4 antibody (BD Biosciences) was used, donor and target cells were preincubated separately with equal volumes of inhibitor for 30 min at 37°C before mixing.

#### Fluorescence microscopy sample preparation

Transfected Jurkat cells (donor cells) were mixed with primary CD4 cells (target cells) in round bottom 96-well-plates for 3-4 hours as previously described. Trim the pipette tips to reduce the shearing to cells. Co-cultured donor and target cells were carefully transferred without disturbance onto poly-lysine treated coverslips. The cells were plated onto the poly-L-lysine treated coverslip for 30 min in 37°C incubator. Media was removed and cells fixed with 4% PFA for 10 min at room temperature, washed twice with PBS, and mounted with anti-fade mounting medium with DAPI (Vectashield, Co#: H-1200, Vector Laboratories). For intracellular staining of Env with 2G12, transfected Jurkat cells were plated onto poly-L-lysine treated cover glass and allowed to attach for 30 min at 37°C. The cells were permeabilized with PBS containing 0.1% triton X-100 and 2% FBS for 5 minutes. Next the cells were stained with 2G12 (1:200) for 1 hour followed by secondary antibody for 45 minutes. After washing, the samples were sealed in mounting media and ready to observe. For surface staining of Env, the cells were directly stained at 4°C with anti-GFP antibody (1:500) diluted in PBS with 2% FBS for 45 min, followed by a secondary antibody for 30 min, and then washed and fixed in 4% PFA or kept alive for live cell pulse-chase experiments.

#### Confocal and live imaging

Confocal imaging was carried out on an inverted Leica SP5 DMI laser scanning confocal microscope, using a 63× objective and analyzed using Volocity (PerkinElmer) or ImageJ (NIH) software. Live imaging was carried out in a sealed, gas permeable microchamber slides (Ibidi Biosciences). Donor cells were mixed with target cells at a ratio of 1:2 and were loaded onto the micro-chamber pre-coated with 150g/ml fibronectin to provide the cells with a two-dimensional substrate for attachment and migration. The chamber was placed on a Zeiss AxioObserver Z1 inverted microscope mounted with Yokogawa CSU-X1 spinning disk scan head. Dual Hamamatsu EM-CCD C9100 digital cameras enable simultaneous imaging of up to two fluorescent channels. Phase contrast imaging and confocal green (for sfGFP) and red (for mCherry) fluorescence were acquired in a multitrack configuration to avoid cross-talk between fluorescence channels. Images were recorded at different time intervals continuously as indicated in results. Confocal images and Quicktime movies were generated from laser-scanning confocal microscope file data using using Metamorph software (Molecular Devices) and Imaris (bitmap) software.

#### Fluorescence Recovery after Photobleaching (FRAP)

FRAP was performed on two systems: Zeiss LSM880 and Leica SP5 DMI. Zeiss LSM880 Airyscan microscope equipped with a 63X oil-immersion objective (NA 1.4) using the 561 nm and 488 nm laser lines. The system is adjusted to proper humidity, 5% CO2 and 37°C. The FRAP experiment on LSM880 used a 4-minute protocol: pre-bleach for 3 sec, bleach for 1 sec at 60% laser power and recovery of fluorescence was captured for the last of the 4 minutes. On Leica SP5 DMI, we used a 60X oil-immersion objective (NA1.4) with 561 nm and 488 nm laser lines. There is an inherent three-step capturing protocol from the system. After 1s bleaching, the first 100 frames were captured continuously; the second 50 frames were at 1s/frame and the last 50 frames at 5s/frame. A rectangular zone covering about half of the virological synapse was bleached, leaving the other half as unbleached area control and localization reference. In one case, where the virological synapse was too small to bleach a fraction of it, a nearby area was selected as unbleached area control. FRAP curve of the bleached virological synapse was determined from ROI rigidly covering the synapse button. A normal bleaching curve was determined from a different area covering most of the cytoplasm of the same cell and used for normalization of values. Fluorescence intensity over time was plotted using GraphPad Prism software, and the data were fitted to a one-phase exponential association function to calculate recovery half-times and immobile fractions.

#### Super-resolution optical microscopy of HIV-infected T cells

3D structured illumination microscopy of fixed T cells cells was performed with a commercial Deltavision OMXv4.0 BLAZE microscope (GE Healthcare, Amersham, UK) using a 60x, 1.42 NA oil immersion PlanApoN objective lens (Olympus, Japan) and sCMOS cameras. Env tagged with sfGFP was excited at 488 nm and the emission recorded at 504–552 nm. Gag tagged with mCherry was excited at 546 nm and the emission recorded at 600-650 nm. The plasma membrane was stained with CellMask Deep Red, excited at 649 nm and the emission recorded at 660-670 nm. The nucleus was stained with DAPI, excited at 405 nm and the emission recorded at 450-470 nm. A sequence of 15 images for each axial plane, obtained at three different angles with five phases each, was acquired. Multiple axial planes encompassing the entire cell from top to bottom were recorded at a separation of the individual axial planes of 125 nm. Super-resolved fluorescent images were reconstructed with the corresponding recorded optical transfer function (OTF) in the SoftWoRx 7.0.0 software (GE Healthcare, Amersham, UK) at a Wiener filter setting of 0.006.

### DATA AND SOFTWARE AVAILABILITY

Primary imaging data are available upon request.

